# EssSubgraph improves performance and generalizability of mammalian essential gene prediction with large networks

**DOI:** 10.1101/2025.07.21.665218

**Authors:** Haimei Wen, Susan Carpenter, Karen McGinnis, Andrew Nelson, Keriayn Smith, Tian Hong

## Abstract

Predicting essential genes is important for understanding the minimal genetic requirements of organisms, identifying disease-associated genes, and discovering potential drug targets. Wet-lab experiments for identifying essential genes are time-consuming and labor-intensive. Although various machine learning methods have been developed for essential gene prediction, both systematic testing with large collections of gene knockout data and rigorous benchmarking for efficient methods are very limited to date. Furthermore, current graph-based approaches require learning the entire gene interaction networks, leading to high computational costs, especially for large-scale networks. To address these issues, we propose EssSubgraph, an inductive representation learning method that integrates graph-structured network data with omics features for training graph neural networks. We used comprehensive lists of human essential genes distilled from the latest collection of knockout datasets for benchmarking. When applied to essential gene prediction with multiple types of biological networks, EssSubgraph achieved superior performance compared to existing graph-based and other models. The performance is more stable than other methods with respect to network structure and gene feature perturbations. Because of its inductive nature, EssSubgraph also enables predicting gene functions using dynamical networks with unseen nodes and it is scalable with respect to network sizes. Finally, EssSubgraph has better performance in cross-species essential gene prediction compared to other methods. Our results show that EssSubgraph effectively combines networks and omics data for accurate essential gene identification while maintaining computational efficiency. The source code and datasets used in this study are freely available at https://github.com/wenmm/EssSubgraph.

## Introduction

A gene is considered essential when the loss of its function compromises viability of the individual (for example, embryonic lethality) or results in profound loss of fitness (Bartha, et al., 2018). The entire complement of essential genes for a given cell type constitutes a minimal gene set for a living cell (Zhang and Lin, 2009). Identification of essential genes in different species not only provides insight into the molecular basis of core biological processes but also sheds light onto potential therapeutic strategies against diseases such as cancer (Kuang, et al., 2021; Yang, et al., 2014). Recently, systematic identification of essential genes in the context of cellular viability has been enabled by new technologies such as genome-wide, CRISPR-based screens (Liu, et al., 2017; Ma, et al., 2015). Nonetheless, experimental identification of essential genes is often expensive, time consuming, and labor intensive. Computational methods can not only provide accurate confirmation of known essential genes but also predict new ones across cell types or species whose essential genes may not have been identified by experiments. These methods can also be useful in predicting functions of poorly characterized genes. However, the performance of computational methods in predicting gene essentiality remains unclear due to the lack of rigorous testing with the latest large-scale collections of experimental data that include hundreds of screens.

Essential genes exhibit distinct patterns across molecular, evolutionary, and developmental dimensions, making these features valuable for predicting gene essentiality (Chen, et al., 2020). In certain cases, genomic and functional data can be used to predict essentiality. For example, Guo et al. used nucleotide composition and internal nucleotide association information to predict essential human genes (Guo, et al., 2017). Kuang et al. used expression data for essential gene prediction (Kuang, et al., 2021). While these methods produce satisfactory results in some contexts, the utility of individual modes of data may be limited with respect to the collective functions with which essential genes support cellular life.

Since genes and their products (proteins or RNAs) interact extensively in cells, essential genes may exhibit distinct patterns within gene or protein interaction networks. This insight allows essential gene prediction to be framed as a node classification problem, where the underlying graph represents a biological interaction network. Network information can be incorporated into machine learning model structures using various methods, such as the factorization-based embedding approach DeepWalk (Perozzi, et al., 2014), Graph Convolutional Network (GCN) (Kipf and Welling, 2017), and Graph Attention Network (GAT) (Veličković, et al., 2017). Some of these network-based approaches have also been applied to predict essential genes. For example, Dai et al. used a Protein-Protein Interaction (PPI) network for human essential genes identification (Dai, et al., 2020). In addition, some recent studies have used graph network-based methods to predict cancer driver genes, including MTGCN (Peng, et al., 2022), HGDC (Zhang, et al., 2023) and EMOGI (Schulte-Sasse, et al., 2021).

PPI network and other information such as sequences can be combined in a deep learning framework for essential gene prediction (e.g. DeepHE (Zhang, et al., 2020)). However, these network-based essential gene prediction methods rely on representation learning with the entire PPI as a starting point of model construction and training, which is neither an efficient approach nor a realistic assumption due to the constant expansion of experimental discoveries and the dynamic nature of biological networks. It is unclear whether accurate predictions of essential genes can be achieved without the prior global information of the PPI network In this study, we developed a method, termed Essential Gene Prediction with Subgraphs (EssSubgraph), that leverages subnetwork sampling with only local network information and expression data to accurately predict essential genes. We show that with widely used gene expression datasets and PPI networks, EssSubgraph not only had significantly better and more stable performance compared to previous models, but also used less prior information, including identity and connectivity of unseen genes. In addition, EssSubgraph has a lower memory requirement than other graph neural network-based approaches, which confers the ability to model large-scale biological networks. The essential genes identified by this model had annotated biological functions as expected from experimentally identified genes. Finally, we applied the model trained with human data to mouse genes and observed satisfactory performance.

## Materials and Methods

### Overview

In this study, we present EssSubgraph, a graph neural network framework based on GraphSAGE (Hamilton, et al., 2017) with new modifications. The model integrates protein-protein interaction (PPI) networks with multi-omics data, including transcriptomic profiles from The Cancer Genome Atlas (TCGA) normalized RNA-Seq data (Bacolla and Tainer, 2023), for essential gene prediction. Specifically, the edges of the PPI networks are underpinned by various types of physical and nonphysical interactions and are obtained from multiple databases. The PPI network structures were obtained from CPDB (Kamburov, et al., 2011), STRING (Szklarczyk, et al., 2023), BioGRID (Stark, et al., 2006), HumanNet (Kim, et al., 2022), IRefIndex (Razick, et al., 2008), PathwayCommons (Rodchenkov, et al., 2020) and PCNet (Huang, et al., 2018). TCGA gene expression data were processed through principal component analysis (PCA) to generate low-dimensional node features. The ground truth labels (essential and non-essential gene lists) for model training were derived from DepMap (https://depmap.org/portal) (Tsherniak, et al., 2017). Our computational framework employs an inductive learning architecture to integrate three key biological data components: (1) PPI networks, (2) dimensionality-reduced gene expression profiles from TCGA (node features), and (3) experimentally validated essential and non-essential gene lists from DepMap (training labels) (**Figure 1**). Through an iterative neighborhood sampling and feature aggregation mechanism, the model derives low-dimensional gene embeddings that encapsulate both network context and functional genomic characteristics. These learned representations subsequently serve as discriminative features for supervised essential gene classification.

**Figure 1:**
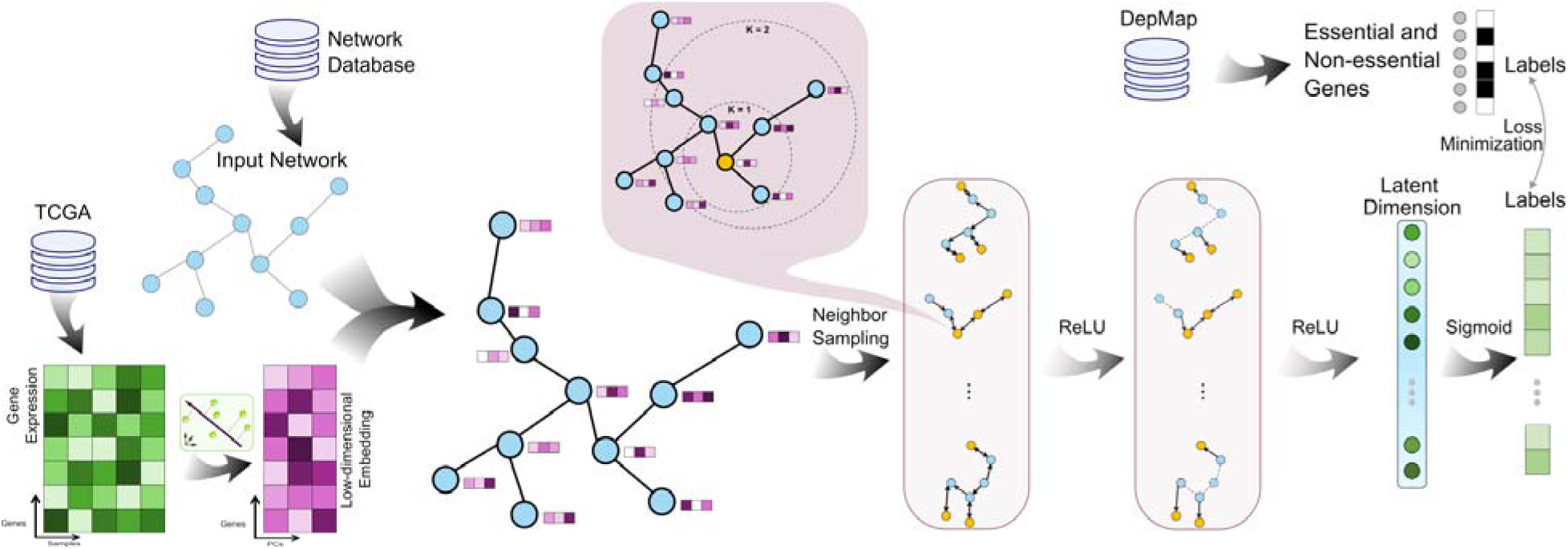
Overview of data sources and EssSubgraph framework for essential gene prediction. Essential genes, non-essential genes were collected from DepMap. Node feature vectors were derived from TCGA expression data with dimensionality reduction. A graph neural network based on PPI network databases is created, trained, and evaluated for essential gene prediction. An aggregation scheme (callout) was used upon random sampling of neighbors for each node.

### Essential Gene Datasets

We obtained 2,299 common essential genes (abbreviated as essential genes) and 10,723 non-essential genes from the DepMap database as positive and negative instances for model construction with a procedure described in this section. Because essential genes are indispensable for most cell types to proliferate, the essential gene detection approach underpinning the DepMap database assumed that if a gene is universally important for cell viability, i.e. if it is a common essential gene, it should produce a significant growth phenotype in the vast majority of cell lines (e.g. >90% in this work) based on high-throughput screening experiments including RNAi and CRISPR (Dempster, et al., 2019). We therefore followed the approach by Dempter et al (Dempster, et al., 2019): We first ranked genes by their gene (viability) effect scores within each cell line, then computed the distribution of these percentile ranks across all cell lines for each gene. We used Gaussian kernel density estimation (KDE) on the minimal percentile ranks to establish an essentiality threshold with the optimal point as the lowest density in the smoothened distribution. Our study on the gene effect data from CRISPR and RNAi revealed that the distribution produced a threshold of around 0.3. Based on this strategy, the common essential genes that we identified was also consistent with the summary from DepMap (https://depmap.org/portal/api/download/gene_dep_summary) using combined CRISPR and RNAi screening data. Therefore, we used the intersection of the common essential genes identified from the DepMap-processed CRISPR and RNAi analyses. Conditionally essential genes were extracted from the DepMap summary file, which were identified through a Likelihood Ratio Test-based method described earlier (Dempster, et al., 2019; McDonald, et al., 2017). The non-essential genes were defined as genes that are neither from the list of common essential genes nor from the list of conditionally essential genes. Note that our approach of selecting common essential genes is similar to previous work (Kuang, et al., 2021) with two key differences: First, we used a much larger dataset containing screening experiments from multiple sources: 1,178 cell lines from CRISPR (DepMap Public 24Q4+Score, Chronos) and 708 cell lines from RNAi (Achilles+DRIVE+Marcotte, DEMETER2); Secondly, our approach considered 6,155 conditionally essential genes as unlabeled samples (nodes) to be included in our model.

To derive the expression-based features for essential, non-essential and unlabeled genes, the TCGA RNA-sequencing data for 20,530 genes in 10,039 cancer samples across 33 cancer types were downloaded from Zencode (Bacolla and Tainer, 2023). The data were normalized in the units of Fragments per Kilobase of transcript per Million mapped reads (FPKM). Details of deriving gene features from expression data are described in a later section.

We obtained PPI networks from CPDB, STRING, BioGRID, HumanNet, IRefIndex, PathwayCommons, and PCNet. We exclusively considered high-confidence interactions in each network with_a___filtering strategy___established earlier (Huang, et al., 2018). For the CPDB network, we kept interactions with a score higher than 0.5, and for STRING we used a threshold of 0.85. After filtering, we obtained 13,261 nodes with 296,428 edges in the CPDB network, 13,137 nodes with 244,978 edges in the STRING network, 20,096 nodes with 865,319 edges in the BioGRID network, 16,190 nodes with 475,867 edges in the HumanNet network, 17,159 nodes with 607,613 edges in the IRefIndex network, 19,087 nodes with 1,040,197 edges in the PathwayCommons network, and 19,781 nodes with 2,724,724 edges in the PCNet network. The graph in each of our neural network models uses only one of these networks. Since the nodes in each network only contain subsets of those labeled and unlabeled genes that we obtained from DepMap (**Supplementary Table S1**), our model focused on training with and predicting those in-network genes for all methods that we compared in this work.

### Model

The structure of the EssSubgraph model is based on GraphSAGE (Hamilton, et al., 2017) with some modifications described in this section. The model is an inductive deep learning method that generates low-dimensional vector representations for nodes in graphs and can predict identities of genes which can then predict the identities of genes that were either included in the training network or are unseen nodes. Therefore, the model does not require the whole network structure during learning and the learned model with node embeddings can generalize to previously unseen nodes (Gao, et al., 2024). In this work, we first follow a method widely used in graph neural networks to investigate the model performance: The training process incorporates the full network topology including all genes, with only the essential labels of test genes being masked to maintain consistency of evaluation. Subsequently, we tested our model with ‘expanding networks’, i.e. the networks at the model training steps only contain the subset of genes used for training, and the testing genes are completely ‘unseen’ before testing.

The model learns node representations by sampling and aggregating neighbors from multiple search depths or hops (i.e. sample and aggregate). Our model first samples or prunes the *K*-hop neighborhood computation graph and then performs the feature aggregation operation on this sampled graph in order to generate the embeddings for a target node. Nodes aggregate information from their local neighbors with an iterative process

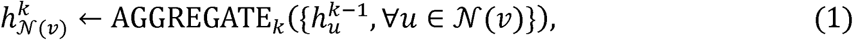

where *v* is the vertex set of the graph, *N(v)* is the vertex set in the immediate neighborhood of a node *v* (∀*v* ∈ *v)*, *k* denotes the current step in the outer loop (or the depth of the search) and *h*^k^ denotes a node’s representation at this step. Each node first aggregates the representations of the nodes in its immediate neighborhood (“base case” *k*= 0). The model then concatenates the node’s current representation, 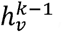, with the aggregated neighborhood vector, 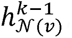 , and this concatenated vector is fed through a fully connected layer with nonlinear activation function *a*, which transforms the representations to be used at the next step of the algorithm. As the iteration proceeds, nodes incrementally gain more information from distant positions in the graph. The mean aggregator is similar to the convolutional propagation rule used in the transductive GCN framework. i.e.

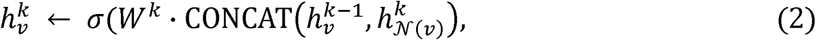

where *W* a learnable weight matrix and *a* denotes a non-linear function (RuLu activation function was used here). To obtain the final, integrated node features for the node classification task, EssSubgraph maps 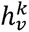 to a low-dimensional space through a learned linear transformation.

For model training, labeled data were randomly split into training (80%), test (20%) through fivefold cross-validation, Additionally, we further split the training set, with 10% used for validation and the remaining for training. We computed the cross-entropy loss *L* for our training node as:

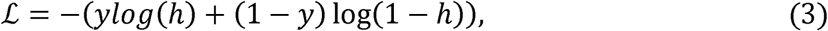

where h is the output of the network after sigmoidal activation layer and *y* the original label (0 or 1). We used torch BCEWithLogitsLoss to implement this functionality. And ADAM optimizer (Shindjalova, et al., 2014) with a learning rate of 0.01 to train the model for 200 epochs. Early stopping was used based on model loss on the validation set.

### Benchmarking

For the evaluation metrics, we used Area Under the Precision-Recall Curve (AUPRC) and Area Under the Receiver Operating Characteristic Curve (AUROC), which is implemented by using sklearn. For benchmarking, 5-fold cross-validations were performed and the mean values AUPRC were used to compare performance. A total of eight previously published methods for essential gene predictions were used to evaluate the performance of EssSubgraph (Cortes and Vapnik, 1995; Kipf and Welling, 2017; Kuang, et al., 2021; Peng, et al., 2022; Perozzi, et al., 2014; Schulte-Sasse, et al., 2021; Veličković, et al., 2017; Zhang, et al., 2020). Among them, four methods were based on graph neural networks. Comparisons on metrics such as memory usage were performed with this group of alternative methods.

### Node features

We used principal component analysis (PCA) to obtain node features from the expression matrix of TCGA. We normalized gene expression data, performed PCA, and Min-Max scaling. To select the optimal number of PCs, we performed a scan with the range of 10-300 PCs. We used the EssSubgraph model mentioned above. With the metrics of the areas under AUROC and AUPRC, we found that 50 PCs performed best among the selected group (**Figure 2A** and **B**). We therefore used the top 50 PCs as the node features for the subsequent tests.

**Figure 2.**
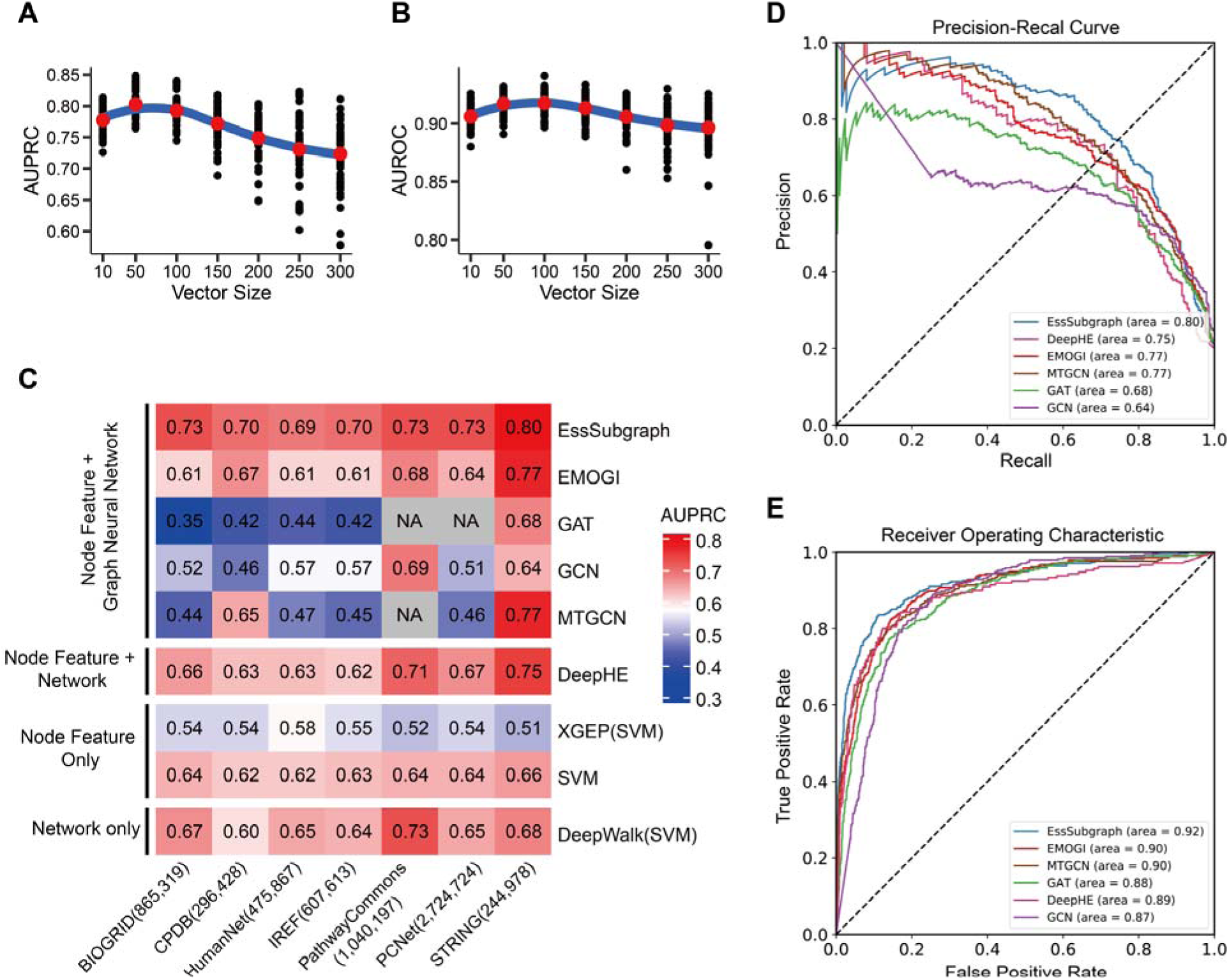
EssSubgraph outperforms previous methods in predicting essential genes. **A**. Effect of Principal component vector size on AUPRC performance. 5-fold cross validations were performed 10 times to ensure stability. **B**. Effect of Principal component vector size on AUROC performance. **C**. Mean AUPRC values from 5-fold cross-validations for different prediction methods across different PPI networks. Dark blue cells in the heatmap correspond to low performance (low AUPRC values), whereas dark red cells correspond to higher performance. Methods are grouped according to the type of data used: network only, methods that only use the PPI network; node feature and network, methods that use both PPI network and transcriptome information; node feature only, methods that only use gene expression feature information. NA indicates that the method did not produce predictions due to high memory cost. The number in parentheses after the network name indicates the number of edges in each network. **D**. Representative AUPRC curves of EssSubgraph and the five other methods. STRING network was used. **E.** Representative ROC curves of EssSubgraph and five other methods (EMOGI, GAT, GCN, MTGCN and DeepHE).

### Gene ontology enrichment analysis of essential genes

Functional enrichment analysis of essential genes was performed using clusterProfiler (version 4.12.6) (Yu, et al., 2012). Gene Ontology (GO) enrichment was restricted to the Biological Process (BP) category. Statistical significance was determined with the Benjamini-Hochberg procedure and a q-value cutoff of 0.05.

### Cross-species predictions

Labels for mouse developmental essential genes were obtained from a published dataset (Kabir, et al., 2017). Mouse gene symbols were converted to human gene symbols using the NicheNet R package (version 2.2.0) (Browaeys, et al., 2020). To demonstrate the model’s ability to be trained for cross-species prediction, human gene expression and network information were used for training as described earlier.

## Results

### Performance comparison with other methods

To evaluate the performance of our model, we used eight benchmark methods, GCN (Kipf and Welling, 2017), GAT (Veličković, et al., 2017), EMOGI (Schulte-Sasse, et al., 2021), DeepWalk (Perozzi, et al., 2014), MTGCN (Peng, et al., 2022), DeepHE (Zhang, et al., 2020), XGEP (Kuang, et al., 2021) and SVM (Cortes and Vapnik, 1995). Among them, XGEP and DeepHE were specifically designed for essential gene prediction but used limited experimental data for evaluation, DeepWalk was used as a network-only benchmark and SVM was used as a network-free benchmark. GCN is a typical graph neural network approach that learns new features by aggregating features from its direct neighbors and itself. MTGCN is a multi-task and multi-graph convolutional network method. XGEP extracts gene features through collaborative learning and subsequently applies classification methods such as SVM, DNN, and XGBoost (Chen and Guestrin, 2016). EMOGI is a recently developed deep learning framework based on graph convolutional networks. It integrates multi-omics (e.g. DNA methylation and gene expression) data as node features and incorporates PPI networks to learn informative gene representations for the prediction of cancer driver genes. DeepHE uses concatenated features from both the graph and sequence, where the graph features are generated using DeepWalk, which performs well when learning from accurate and relatively small graph datasets. Among these benchmark methods, Graph Neural Networks (GNNs) offer an advantage by more comprehensively capturing and reflecting the structural information of the network (**Figure 2C** top). In each test for network-based predictions, we used one PPI network and the TCGA-derived gene expression (node feature) matrix in each method for comparison. We tested multiple PPI networks from different databases with varying network complexity.

We applied EssSubgraph and alternative methods to predict essential human genes (2,299 essential genes and 10,723 non-essential genes) and used AUPRC as the main evaluation metric because it is suitable for class-imbalance prediction. We also showed AUROC as a supporting metric. EssSubgraph achieved satisfactory performance with the STRING network (AUPRC 0.80) and other PPI networks. Across all networks, EssSubgraph had a better performance than each of the benchmark methods for essential gene prediction (**Figure 2C and D**). This result suggests its effectiveness in learning from various types of local topology and generalizes well to new structures.

Some earlier studies reported higher values of AUPRC with lists of essential genes from older datasets. For example, Kuang et al. used a list of 1516 essential genes selected from screening experiments with 11 cancer cell lines (Guo, et al., 2017; Kuang, et al., 2021) and achieved AUPRC 0.83. Unsurprisingly, this gene list is more restricted than the one used in this work generated from a large collection of experiments. To test the performance of different methods with this older dataset, we used a previously published list of essential and non-essential genes (Guo, et al., 2017; Kuang, et al., 2021) to construct the labels and compared the performance of EssSubgraph with EMOGI, MTGCN, GCN, GAT, and SVM, using AUROC and AUPRC as evaluation metrics. EssSubgraph again achieved the best performance (AUPRC 0.90), followed by DeepHE, MTGCN and other methods (see **Supplementary File Table S2**).

### Expression patterns and network topology contribute to model performance

Since we used both transcriptome and network topology data in essential gene prediction, we next asked whether the performance depends on the combination of these two data types. We performed several perturbations on the edges in the STRING network, the feature vectors of individual genes, or both at the same time, and evaluated the performance. Edge perturbation was performed with random selection of vertices from the network. We scanned the proportions of perturbed edges in a range from 0 (no perturbation) to 100% (all edges were perturbed) (**Figure 3** brown). As expected, we observed that the perturbed networks became dissimilar to scale-free networks (**Supplementary Figure 1**). Similarly, for node feature perturbations, we permuted the feature vectors between pairs of nodes for 25%, 50%, 75%, and 100% of the nodes in the network (**Figure 3** pink). Finally, we perturbed 25%, 50%, 75% and 100% of both nodes and edges (**Figure 3** purple).

**Figure 3.**
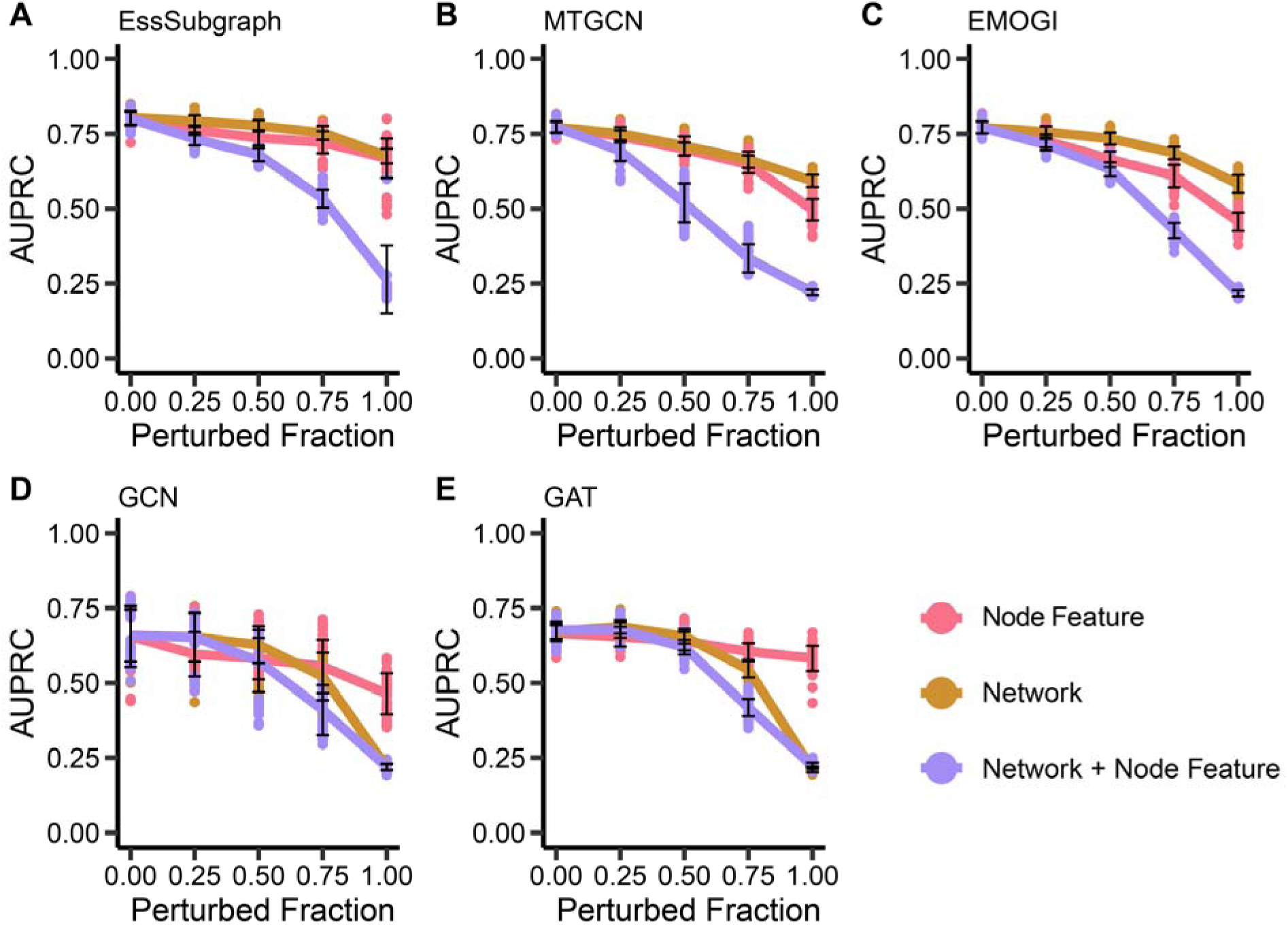
Performance of different methods upon perturbation of node features and network information. **A**. Test set performance of EssSubgraph after systematic perturbation of node features and network information. **B**. Test set performance of MTGCN after systematic perturbation of node features and network information. **C.** Test set performance of EMOGI after systematic perturbation of node features and network information. **D**. Test set performance of GCN after systematic perturbation of node features and network information. **E**. Test set performance of GAT after systematic perturbation of node features and network information.

We found that perturbing both node features and edges significantly decreased the AUPRC values for EssSubgraph as well as four other graph neural network-based methods (**Figure 3**). For EssSubgraph and MTGCN, the performance exhibits moderate decrease when either node feature or network structure was perturbed (**Figure 3A and 3B**). For EMOGI, performance decreased more significantly when perturbing only node feature compared to perturbing solely the network structure (**Figure 3C**). The performance of the GCN and GAT models seems to be very sensitive to large perturbation of network structures (**Figure 3D and E**). Overall, EssSubgraph had a satisfactory performance with moderate perturbations of node features and networks. Furthermore, the large decrease of performance with the combined perturbation of both types of data compared to individual types suggest that these two types of information have the ability to compensate for the loss of each other within our model framework, even though their data structures and their sources are different.

### Memory efficiency and network scalability of EssSubgraph

Some realistic biological networks have high density, such as PCNet (more than two million edges and 138 edges per node) (Huang, et al., 2018). When we tested various methods with a widely used graphics processing unit (GPU), Nvidia 2080 Ti with 12-gigabytes (12GB) of memory, we found that the usage of graph modeling methods such as GCN and GAT exceeded the memory capacity of the GPU and failed to perform training with this type of large network. To examine the dependency of the usability of different methods on network sizes more systematically, we simulated a series of networks with 5 node numbers ranging from 10,000 to 30,000, and 9 edge numbers from 200,000 to 1,800,000, representing 45 networks with various sizes and densities (**Figure 4**).

**Figure 4.**
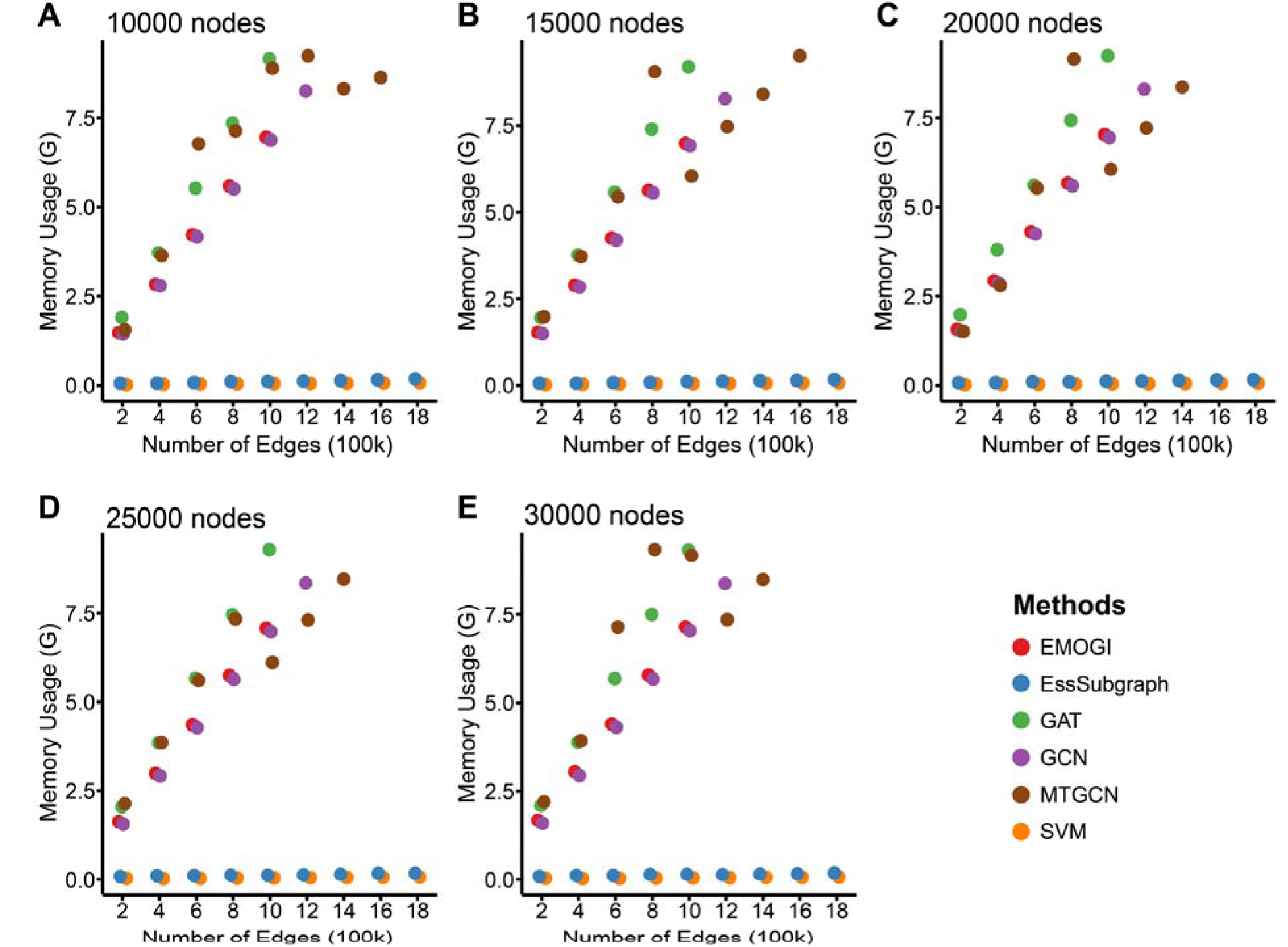
Comparison of memory usage with varying network sizes. Memory usage (GB) of varying node numbers (10,000 to 30,000) with edge numbers (in 100,000). Each subplot represents a fixed node number and multiple numbers of edges. Different colors correspond to different methods: EMOGI (red), GAT (green), GCN (green), EssSubgraph (blue), and SVM (orange). The results show that memory usage increases with the number of edges, with variations depending on the method used. Missing dots in each plot indicates the failure of methods in training models due to high memory usages.

We found that for networks with the same number of nodes and edges, the GAT and MTGCN methods require the most memory, followed by EMOGI and GCN (**Figure 4**). This is likely due to GAT’s attention mechanism, which computes pairwise attention scores between nodes, leading to increased memory consumption. Missing circles in **Figure 4** with some models indicate that these models exceeded the GPU memory capacity of the workstation and failed to produce results. This highlights the scalability limitations of certain architectures. In contrast, EssSubgraph and SVM (SVM is a reference model without network information) have the lowest GPU memory requirements (**Figure 4** blue and orange circles), suggesting that EssSubgraph is suitable for larger networks or resource-constrained environments. Moreover, since the sampled nodes in EssSubgraph are predetermined, the runtime remains essentially fixed regardless of the overall network size. In contrast, other methods require increasing computations as the network size grows due to their dependency on global adjacency relationships and computations over the entire graph. Therefore, EssSubgraph is a scalable approach in scenarios where computational efficiency and memory constraints are critical.

### Essential genes predicted by EssSubgraph have the same biological function as those within the ground truth data

Essential genes are crucial for an organism’s survival, with their essentiality primarily determined by their biological functions. We next asked whether predicted essential genes have similar functions to experimentally identified ones. We used the ROC curve to identify the decision boundary that maximizes the true positive rate (TPR) while minimizing the false positive rate (FPR). We used the ROC curve to identify the optimal decision boundary by selecting the point that maximizes the Youden’s J statistic (Youden, 1950) (J = TPR − FPR), which represents the best trade-off between sensitivity and specificity. This turning point corresponds to the threshold where the difference between the true positive rate and the false positive rate is greatest, ensuring a balanced and effective classification. According to this criterion, we selected a prediction probability cutoff of 0.1954 for identifying essential genes (**Supplementary Figure 2A**), which resulted in 3,521 predicted essential genes, among which 1,881 overlapped with the ground truth set of essential genes (**Supplementary Figure 2B**).

We performed an enrichment analysis to examine the annotated functions of these essential genes. Given the high performance of the EssSubgraph model, we focused on the annotations provided by Gene Ontology (GO) database for these predicted essential genes. We first performed the GO enrichment analysis on experimentally validated essential genes (i.e. the ground truth) and found that these genes are mainly enriched for ribonucleoprotein complex biogenesis and RNA splicing functions (**Figure 5A**). We performed GO enrichment on predicted essential genes and found similar results. **(Figure 5B**). We performed an overlap analysis of Biological Process (BP) terms derived from GO enrichment (with p-value cutoff = 0.01 and q-value cutoff = 0.05) between the ground truth and predicted essential genes. The ground truth genes were enriched in 601 BP terms, while the predicted essential genes were enriched in 888 BP terms, among which 589 terms overlapped, accounting for 98.0% of the terms enriched by the ground truth essential genes (**Figure 5C**).

**Figure 5.**
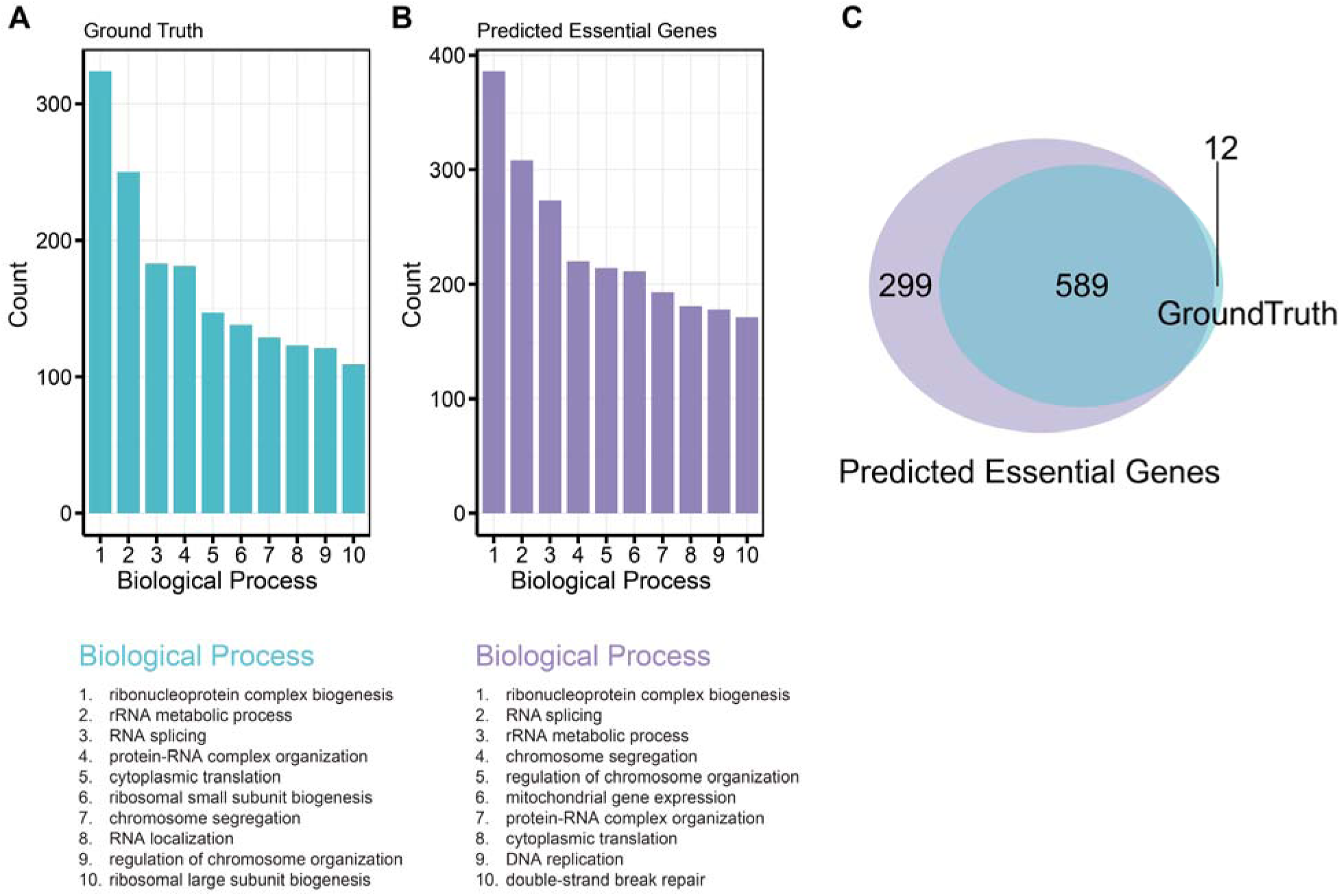
Distributions of top GO terms with validated and predicted essential genes. **A**. Top 10 biological process terms for ground truth essential genes. **B**. Top 10 biological process terms for Esssubgraph predicted essential genes. **C**. The overlap of enriched Biological Process (BP) terms between ground truth essential genes and predicted essential genes. The ground truth set was enriched in 601 BP terms, while the predicted essential gene set was enriched in 888 terms. Notably, 589 BP terms were shared between the two sets, representing 98.0% of the enriched terms in the ground truth set.

### Predicting Essentiality of Unseen Genes

Subgraph neighbor sampling in EssSubgraph offers a key advantage in handling unseen nodes. Unlike traditional full-graph methods such as GCN and GAT, which aggregate information from all nodes, subgraph-based sampling can exclude test and/or unseen node information from the training process while still achieving strong performance. This makes it particularly suitable for scenarios where the network continues to grow (e.g. the expansion of knowledge of gene regulation), and the ability to predict unseen nodes/genes without additional training makes the method more efficient and less prone to errors. We therefore applied EssSubgraph to dynamic biological networks with an approach similar to the application of subgraph methods in other fields (Huang, et al., 2023).

We first used the earliest available version of the STRING network, v9, which was downloaded for training. The filtered STRING v9 version contains 236,440 edges and 9,610 nodes. The trained model was then tested with the latest version, v12 (13,137 nodes with 244,978 edges). The average AUPRC value was 0.80, a performance comparable to the model trained with the latest full network.

Next, we asked how the complete removal of training gene features (network and/or node) from the training process can affect performance. We used three types of perturbations (**Figure 6**) to our original model with the STRING network (**Figure 2D and E**): 1. We removed the test genes from the graph but kept the gene expression matrix of all genes in the dimensionality reduction (PCA) for extracting node features; 2. We retained all nodes of the graph, but we obtained the PCs only with the training nodes and then used it to project all genes, including the unseen ones during testing; and 3. We removed the test nodes completely for training, and added them back to the graph and the PC projection only at the testing step. The AUPRC values obtained for these three different treatments were 0.79, 0.67, and 0.66, respectively. This shows that the lack of network information of the test genes at the training stage had a noticeable but limited impact on the performance of the model. While the gene expression distribution of test genes contributed to the model training significantly, EssSubgraph without this type of data still yielded satisfactory performance comparable to most other methods that leveraged this information (**Figure 2C**). This demonstrates that when node features remain unchanged, applying pre-trained models on evolved network structures alone can still achieve reasonable performance, while the training process itself is computationally expensive.

**Figure 6.**
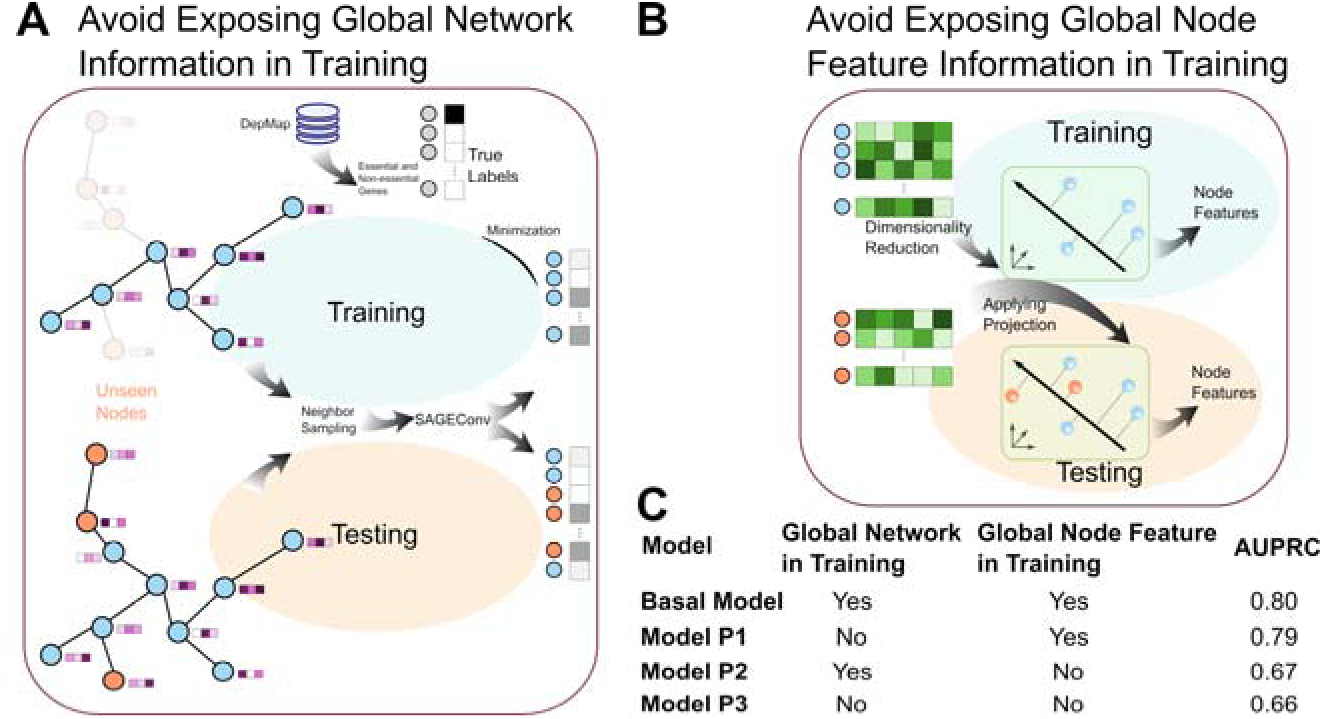
Predictions of unseen genes. **A**. Illustration of training without network information of genes (nodes) to be used for testing. **B**. Illustration of training without expression information of genes (nodes) to be used for testing. **C**. Performance comparison of four models with various combinations of information used for training.

### Cross-species prediction of essential genes

We next investigated EssSubgraph’s applicability across species by using the model to predict essential genes in mice. We used network and node features from human as described earlier and a list of mouse essential genes and non-essential genes from a previous publication (see method) to train and test models. AUROC and AUPRC were again used to assess the classification performance of EssSubgraph and four other graph-based methods. With an AUROC of 0.79 and an AUPRC of 0.49, EssSubgraph outperformed EMOGI, MTGCN, and GCN (**Supplementary Figure 3**), suggesting that the method is suitable for making predictions of essential genes in species or under new conditions where the experimental data may be scarce or unavailable.

## Discussion

Essential genes encode proteins and enzymes that are indispensable for fundamental cellular processes, including maintenance of homeostasis, growth, and development. Impairment or loss of these genes may result in the inability of the organism to survive (Liang, et al., 2024). Identifying essential genes is crucial for elucidating the core cellular architecture and functions, facilitating drug target discovery, guiding synthetic organism design, defining the minimal requirements for cellular life, and uncovering genotype–phenotype relationships (Xu, et al., 2024). Predicting essential genes lays the foundation for many types of gene function identification. In this study, we used the topology of PPI networks and expression data to predict mammalian essential genes. We showed that EssSubgraph, a sub-network sampling approach in a graph neural network framework, improved predictions compared to previous methods. EssSubgraph is based on inductive learning, in which the model can only see the training data and is trained through a sample partial graph or a set of subgraphs. Thus, the generated model and graph embeddings can be generalized to unseen nodes (Gao, et al., 2024). The high performance of this essential gene prediction method complements time-intensive, whole-genome knockout experiments that screen essential genes. We showed that our approach can predict genes whose network and expression features are completely unexposed to the model during training. This is a more realistic scenario than commonly adopted procedures in network-based machine learning approaches, which only avoid the exposure of labels (essential vs non-essential identities), but not gene features to the model. Our approach therefore uses less prior information for model training. This advantage increases the transferability of our model to predicting functions of newly discovered genes or genes that are newly incorporated into the network without re-training the model.

This work focuses on networks of protein-coding genes for which knockout phenotypes are well established experimentally. One important expansion of gene interaction networks is the inclusion of non-coding RNAs (ncRNAs). While limiting our scope to protein-coding genes in this work helped to benchmark multiple methods carefully, our method can be used to predict functions of ncRNAs whose functions are poorly characterized in general. In particular, the genomes of many species, such as humans, have a massive number of long non-coding RNAs (lncRNAs) which play a myriad of roles at cellular and organismal levels (Statello, et al., 2021). Due to the poor sequence conservation of lncRNAs (Sarropoulos, et al., 2019), it is difficult to predict their functions based on sequence information alone. Our work provides a potential approach of predicting lncRNA functions across species based on their homology or synteny arising from their placements in gene interaction networks that contain both protein-coding and ncRNA genes. Nonetheless, future work is needed to establish the benchmark for predicting ncRNA functions which have limited documentation compared to protein-coding genes.

While we did not consider other modes of data, such as sequence features (Guo, et al., 2017; Zhang, et al., 2020), in this study, we demonstrated prominent synergy between node (gene) features and the network (graph) structures for supporting predictions in our framework. This method can be therefore applied to multimodal integration, including other modes of omics data and sequence features, in future work of gene function prediction.

We demonstrated the low memory cost of EssSubgraph, which is nearly the same as models without network information. An advantage of subnetwork sampling over other graph neural networks was also observed by Seon et al. (Seon, et al., 2023). With the expansion of knowledge about gene products, the numbers of nodes and edges can increase significantly, which can raise the requirement of memory (particularly GPU memory) for training models. As the computational power offered by GPU is leveraged more widely in biological research communities (Dematté and Prandi, 2010), the advantage of using sub-network sampling which offers greater GPU efficiency for large-scale networks can become more useful in real-world applications where GPU resources are shared by multiple researchers.

We expect that the EssSubgraph framework proposed here can be used to integrate large-scale omics data and as well as multiple types of networks beyond the ones from individual sources used in this work. For example, this method can be used to predict gene functions in many data-rich applications such as predicting cancer driver genes. Our method provides an important analysis tool for future work in both fundamental research in biology and precision medicine.

## Code Availability

Computer code for reproducing results and applications with other datasets is available at the GitHub repository for this manuscript https://github.com/wenmm/EssSubgraph.

