## Supplementary for "EssSubgraph improves performance and generalizability of mammalian essential gene prediction with large networks"

### Supplementary Figures

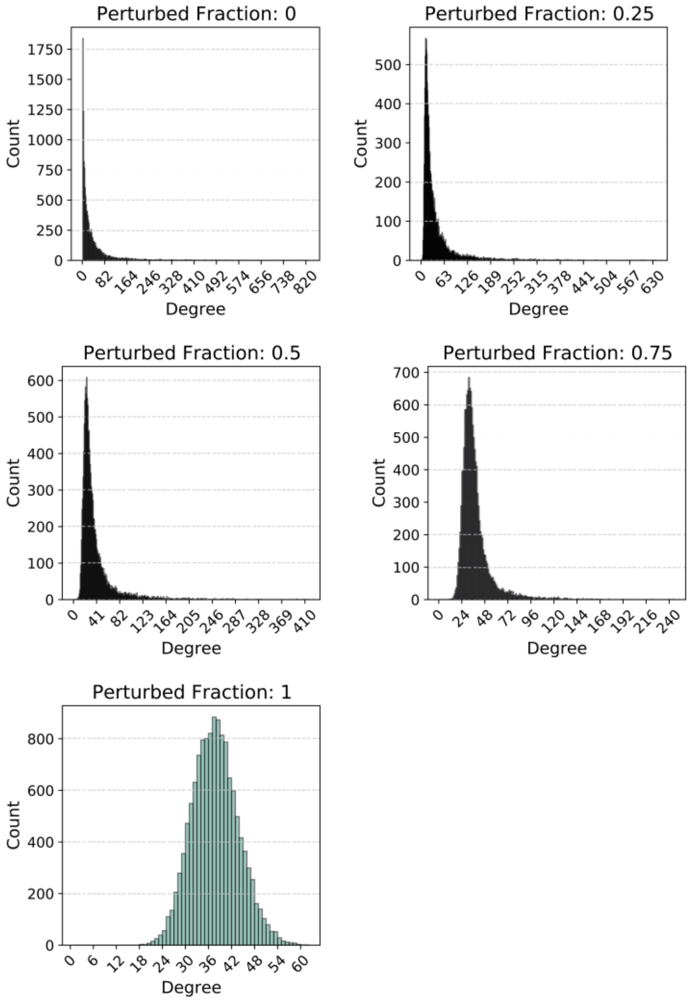


**Figure S1. Degree distributions of the STRING network with perturbations**. Human STRING network of PPI was perturbed by randomly selecting vertices for the indicated fraction of edges.

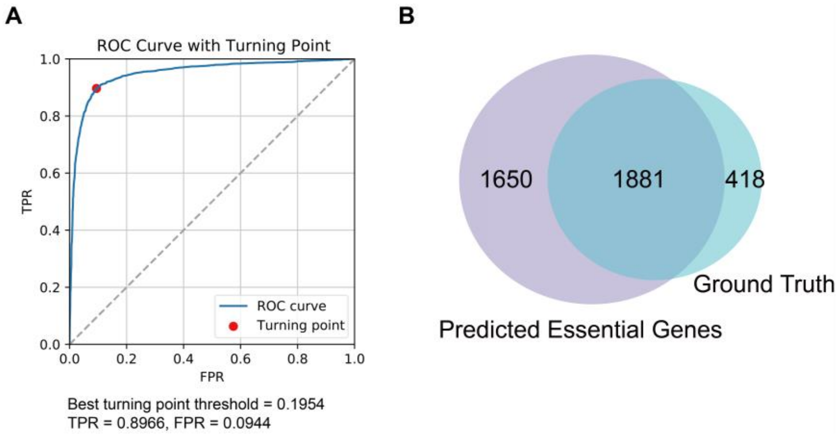


**Figure S2**. **Selection of a model for predicting essential genes**. **A**. The turning point on the ROC curve for selecting the decision boundary. **B**. Overlap between predicted essential genes and experimentally validated ones.

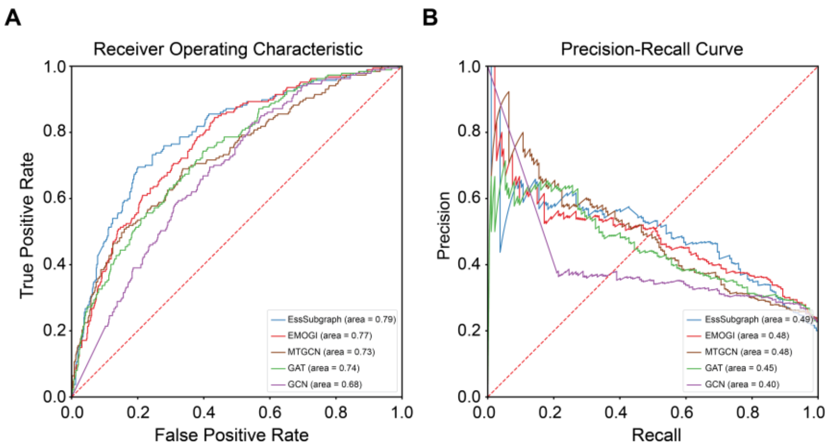


**Figure S3**. **Prediction of mouse essential genes.** A model was trained with a human network and expression data (Figure 2) and was used to predict essential mouse genes. Similar models were used with benchmark methods. **A**. Representative AUROC curves for EssSubgraph and five other graph neural network-based models (EMOGI, GAT, GCN, MTGCN). **B**. Representative AUPRC curves for benchmark models.

### Supplementary Tables

#### Table S1. Gene counts of DepMap and network databases

| Source | Total | Common essential | Unlabeled (conditionally essential) | Non-essential | Unlabeled (others) |
|---|---|---|---|---|---|
| DepMap | 19177 | 2299 | 6155 | 10723 | 0 |
| STRING | 13137 | 2099 | 4452 | 6353 | 233 |
| BIOGRID | 20096 | 2130 | 5682 | 9248 | 3036 |
| CPDB | 13261 | 1975 | 4604 | 6605 | 77 |
| HumanNet | 16190 | 2188 | 5415 | 8326 | 261 |
| IREF | 17159 | 2085 | 5544 | 9114 | 416 |
| PathwayCommons | 19087 | 2202 | 5883 | 9690 | 1312 |
| PCNet | 19781 | 2178 | 5884 | 9715 | 2004 |

**Table S2. Performance of models with labels from Guo et al. 2017**

| Method | AUROC | AUPRC |
|---|---|---|
| EssSubgraph | 0.9715 | 0.8976 |
| EMOGI | 0.9510 | 0.8464 |
| MTGCN | 0.9531 | 0.8585 |
| GCN | 0.9211 | 0.7815 |
| GAT | 0.9224 | 0.7291 |
| XGEP(SVM) | 0.9144 | 0.8285 |
| DeepHE | 0.9559 | 0.8586 |
| SVM | 0.8252 | 0.7511 |
